# Direct Binding of Cysteine-367 Thiolate to the Active Site of the [FeFe]-Hydrogenase from *Clostridium beijerinckii* in the O_2_-stable State

**DOI:** 10.64898/2026.07.11.737921

**Authors:** Jifu Duan, Federica Arrigoni, Andreas Rutz, Eckhard Hofmann, Claudio Greco, Thomas Happe

**Author notes:** **Corresponding Authors** Jifu Duan – Photobiotechnology, Faculty of Biology and Biotechnology, Ruhr University Bochum, 44801 Bochum, Germany;, Thomas Happe – Photobiotechnology, Faculty of Biology and Biotechnology, Ruhr University Bochum, 44801 Bochum, Germany. These authors contributed equally (Jifu Duan, Federica Arrigoni).

## Abstract

[FeFe]-hydrogenases are very active biocatalysts for H_2_ conversion. However, their active site is vulnerable to irreversible degradation initiated by O_2_ binding at the catalytic iron ion (Fe_d_) of the active center. CbA5H, the [FeFe]-hydrogenases from *Clostridium beijerinckii* exhibits stability towards oxygen (O_2_) due to its ability to reversibly enter an inactive state termed H_inact_ upon contact with O_2_. We previously proposed that the close distance of approximately 3.1 Å between the thiol of a nearby cysteine (C367) and the Fe_d_, based on a 2.9 Å crystal structure of CbA5H in the H_inact_ state, enables their binding to each other. This binding therefore was suggested to shield the Fe_d_ from O_2_ damage. However, there is currently a lack of evidence to support this hypothesis. Furthermore, density functional theory (DFT) calculations based on a homologous model favored hydroxide as the binding ligand of the Fe_d_ over the thiol of C367. In this study, we present the crystal structure of CbA5H in the H_inact_ state at an improved resolution of 2.15 Å. The structure reveals a direct binding between the thiol of C367 and the Fe_d_ with a distance of approximated 2.77 Å which is well supported by our DFT calculations based on the new crystallographic data. It is noteworthy that the 2.77 Å bond distance is strikingly long when compared with other iron-sulfur bonds. This finding may provide a crucial foundation for understanding the rapid reversibility of the H_inact_ state.

## INTRODUCTION

[FeFe]-hydrogenases catalyze reversible interconversions between protons, electrons and molecular hydrogen (H_2_) with turnover frequence of up to 10,000 s^-1^,^*1*^ representing the most efficient H_2_ biocatalysts. This extraordinary efficiency is enabled by the uniquely structured hexanuclear iron-complex termed H-cluster and a well-tuned protein scaffold.^*2*^ The H-cluster is composed of a canonical [4Fe4S] subcluster ([4Fe]_H_) and a diiron moiety ([2Fe]_H_) which are covalently connected by a thiolate of a cysteine from the protein environment (Figure 1).^*3, 4*^ Depending on proximal and distal positions relative to the [4Fe]_H_, the two iron ions of the [2Fe]_H_ are denoted as Fe_p_ and Fe_d_, respectively. The Fe_p_ and Fe_d_ ions are coordinated by carbon monoxide (CO) and cyanide CN^−^ ligands.^*5*^ Besides a shared CO ligand in the bridging position, the two iron ions are further connected via an azadithiolate ligand (ADT).^*6-8*^ This configuration creates a vacant site at the Fe_d_ for the binding of substrates^*9-12*^ and inhibitors^*13-16*^ (Figure 1A). The amine group of the ADT ligand exchanges protons with the solvent through a conserved proton transfer pathway (PTP) for catalytic turnover.^*17-20*^

**Figure 1.**
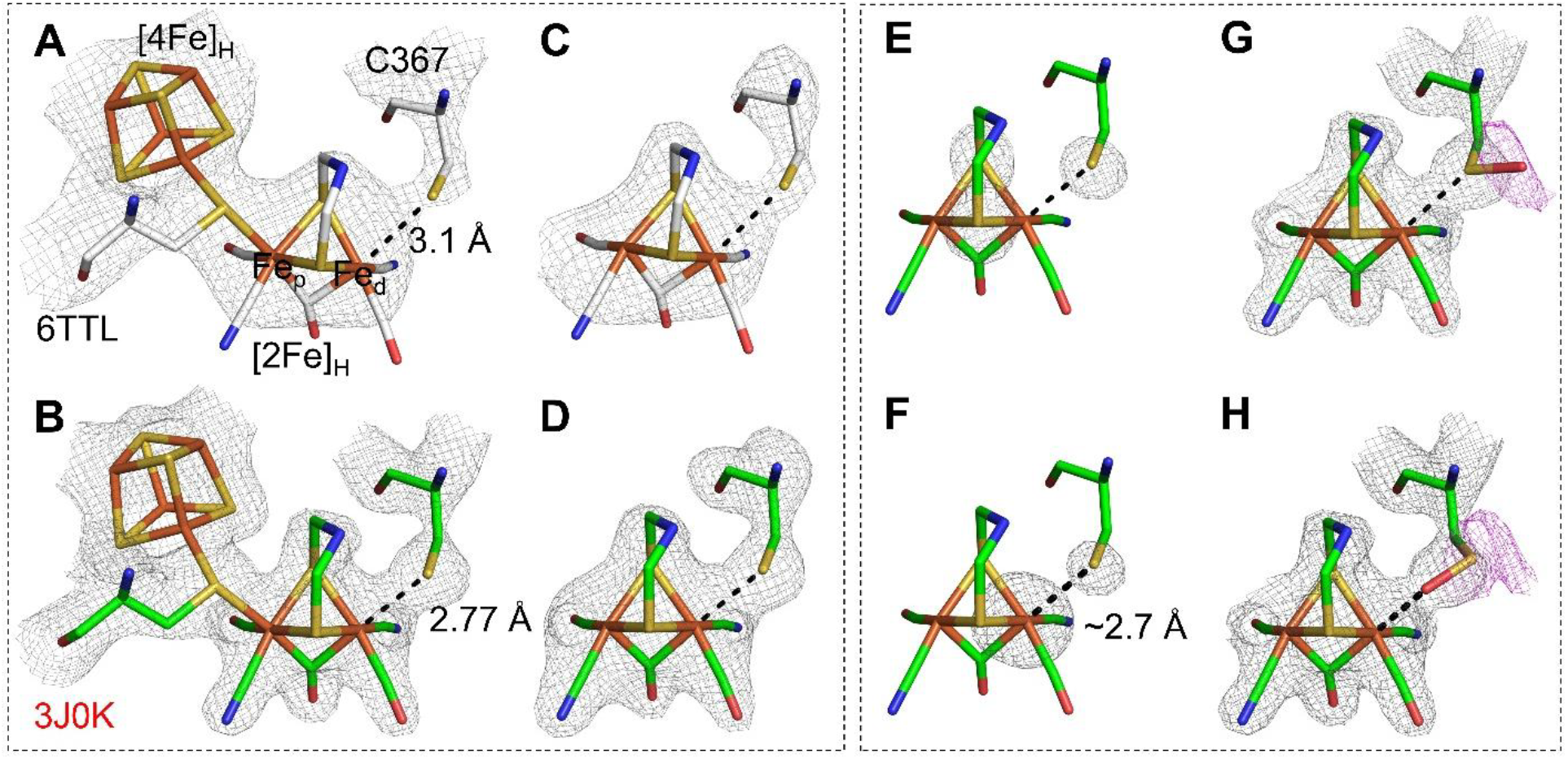
Electron density maps of the H-cluster and C367 of CbA5H in the H_inact_ state. **A** and **B** show 2mFo-DFc maps of the H-cluster and C367 of CbA5H in the H_inact_ state from structures of 6TTL and the new structures presented here (PDB: 30JK). Panels **A-D** show comparison of density maps of the previous structure (6TLL; **A** and **C**) and the new structure (30JK; **B** and **D**) from this study with a focus on the H-cluster and C367. The two structures are distinguished by coloring carbon atoms white and green for 6TTL (**A, C**) and 30JK (**B, D, E-H**), respectively. **E** shows densities of three sulfur atoms, namely two from the dithiolate of the [2Fe]_H_ and one from the thiol of C367. **F** shows individually calculated omitting maps of the Fe_d_ of the [2Fe]_H_ and thiol of C367. **G**-**H** show two modeling schemes of sulfenic acid at position 367. 2mFo-DFc maps and mFo-DFc omitting maps were colored gray and shown in **A/B**/**G**/**H** and **C**/**D**/**G**/**H**, respectively. Additional negative mFo-DFc maps colored magenta are shown in **G** and **H**. The contour levels of the maps are 2.5 σ in **A**/**B**, 3.2 σ in **C**, 4.5 σ in **D**, 7σ in **E**, 9 σ in **F** and 2.5 σ in **G**/**H**. The distances between the Fe_d_ and the thiol of C367 are indicated in **A**/**B**/**F**. The figure is prepared based on structural information from chain A. Results obtained from chain B are presented in Figure S1.

Despite their high activity, most [FeFe]-hydrogenases are very sensitive to molecular oxygen (O_2_). Their H-clusters are subject to irreversible degradation initiated by O_2_ binding at the vacant site of the Fe_d_.^*21-25*^ [FeFe]-hydrogenase from *Clostridium beijerinkii* termed CbA5H is one of several examples exhibiting remarkable resistance to O_2_. Its H-cluster transiently forms an inactive but protected state (H_inact_) in the presence of O_2_.^*26-29*^ H_inact_ can also be formed in a few other [FeFe]-hydrogenases, for example DdH from *Desulfovibrio desulfuricans*^*1, 14, 30, 31*^, ToHydA from *Thermosediminibacter oceani*^*32*^ and CpIII from *Clostridium pasteurianum*^*33, 34*^. The formation of H_inact_ requires oxidation of the H-cluster from the active states. Notably this oxidation does not necessarily result from O_2_ because treatment with other oxidants under anoxic conditions also triggers the formation of H_inact_^*26, 31*^. Additionally, the formation of H_inact_ in DdH is dependent on the presence of sulfide. It has been demonstrated that sulfide directly binds to the Fe_d_ in DdH to form the H_inact_ state so that the [2Fe]_H_ can be protected against O_2_.^*14, 31, 35*^. In contrast, the formation of H_inact_ in CbA5H, ToHydA and CpIII does not depend on extrinsic sulfide and is a built-in property instead.^*26-28, 32-34*^ Besides oxidation of the H-cluster, transition from the active states to the H_inact_ state in CbA5H is accompanied by a conformational rearrangement of a secondary loop close to the H-cluster. This results in a distance change between the Fe_d_ and the thiol of proton-transfer C367 on the loop from about 6 Å to 3.1 Å, as shown in structures of CbA5H in the active and H_inact_ states resolved at 2.2 and 2.9 Å, respectively.^*28, 29*^ Based on the relatively close distance between the Fe_d_ and the thiol of C367 in the H_inact_ state, we postulated that it is a direct binding between them. This hypothesis was supported by site-directed mutagenesis experiments. Exchanging C367 to alanine and aspartate largely abolished the formation of H_inact_. However, density functional theory (DFT) calculations favor a hydroxide over the thiol group as the ligand of the Fe_d_.^*27*^ In order to determine the actual structure of H_inact_ in CbA5H, higher resolution structural data are essential.

## RESULTS AND DISCUSSION

Here, we improved the resolution of the structure of CbA5H in the H_inact_ state to 2.15 Å crystallized under aerobic conditions (see details in method part). The overall architecture is very similar to the previous structure. The asymmetric unit contains two copies of a protomer, namely chains A and B from the known CbA5H homodimeric structure.^*29*^ The higher resolution structure shows electron densities with significant improvement. Taking the H-cluster region as an example, the previous structure only showed two big, undefined regions of densities from the two subclusters, whereas the profile of the H-cluster including the CO and CN^−^ ligands is clearly visible in the improved structure (Figure 1A-D and Figure S1A-D). The electron densities of C367 including the thiol group are well-defined, too. The thiol group of C367 shows comparable electron densities with those from the two sulfur atoms of the [2Fe]_H_, suggesting consistent rigidness of the H-cluster and C367 (Figure 1E). The two centers of the omit maps of the Fe_d_ and the thiol of C367 are estimated to be about 2.7 Å apart (Figure 1F). Modeling a hydroxide between these two atoms resulted in strong clashes between the hydroxide and the thiol of C367 with the distances of 2.17-2.24 Å, much higher temperature factors of the hydroxide than other surrounding atoms, no meaningful densities of the modeled hydroxide ion and strong unexplained densities of C367 (Figure S2). Thus, these structural results strongly suggest that the presence of a hydroxide as a protecting ligand of Fe_d_ in the H_inact_ state of CbA5H is highly unlikely. Although the oxidation of C367 to sulfenic acid (-S-OH) does not appear to be relevant for the formation of H_inact_ in CbA5H,^*27, 36*^ we still modeled this species for several reasons. First, this species had a better fit than the thiol of C367 in previous DFT calculations.^*27*^ Second, the oxygen atom of sulfenic acid would result in similar electron densities as that of a hydroxide anion, modeling sulfenic acid may enable us to gain insights into the fit of an additional oxygen atom in this region when this oxygen atom is covalently bound to the sulfur atom of C367. Two sulfenic acid modeling schemes were obtained depending on the nature of the atom coordinating to the Fe_d_, that is the sulfur or the oxygen of the sulfenic acid at position 367. In both cases, strong unexplained negative densities were found around the oxygen or sulfur atom of the modeled sulfenic acid (Figure 1G-H). These densities are absent when a cysteine was modeled (Figure 1B-D), suggesting a much better fit with cysteine than sulfenic acid. Hence, the improved structure strongly suggests the thiol of C367 as the ligand that binds to the Fe_d_ in the H_inact_ state of CbA5H.

As indicated above, this distance between the Fe_d_ and the thiol of C367 was estimated to be about 2.7 Å, which is much longer than the common iron-sulfur bond length of about 2.3 Å.^*37*^ Therefore, we performed systematic and interactive refinements to determine this bond length more precisely. The first refinement cycle was performed with a strict restraint (standard deviation σ of 0.005 Å) of this bond distance of 2.3 Å. The resulting bond distances in chains A and B from the first cycle of refinement were 2.3 Å (Figure 2), as expected. However, significant negative electron densities between the Fe_d_ and the thiol of C367, and simultaneous significant positive electron densities locating on the other side of the thiol of C367 were found (Figure S3), strongly suggesting the length restraint resulted in the thiol of C367 being too close to the Fe_d_. In the next cycle of refinement, when the bond distance restraint was relaxed with σ value of 0.1 Å, the resulting bond distances increased to about 2.45 Å. Applying the same relaxed restraint for the bond distance between the Fe_d_ and thiol of C367, seven iterative refinement cycles were performed until the bond distances remained unchanged (with deviations of less than 0.02 Å compared to the results from a previous refinement cycle), resulting in distances of 2.77 and 2.7 Å in chains A and B, respectively. Similarly, the interactive refinement was performed with a strict restraint of the bond distance to 3.1 Å. The resulting maps from cycle 1 suggested that the distance is too large because positive electron densities are now apparent between the thiol group of C367 and Fe_d_ (Figure S3E-F). After relaxing the restraint in the following cycle as done above, the refined distances dropped to 3.02 and 2.96 Å in chains A and B, respectively (Figure 2). Following six interactive refinement cycles with the relaxed restraint, the bond distances gradually decreased until they remained unchanged with the final distances of 2.85 and 2.77 Å in chains A and B, respectively. These values are very similar to the values obtained by interactive refinements starting with 2.3 Å described above. By performing these systematic refinements, an averaged distance of 2.77±0.06 Å was obtained.

**Figure 2.**
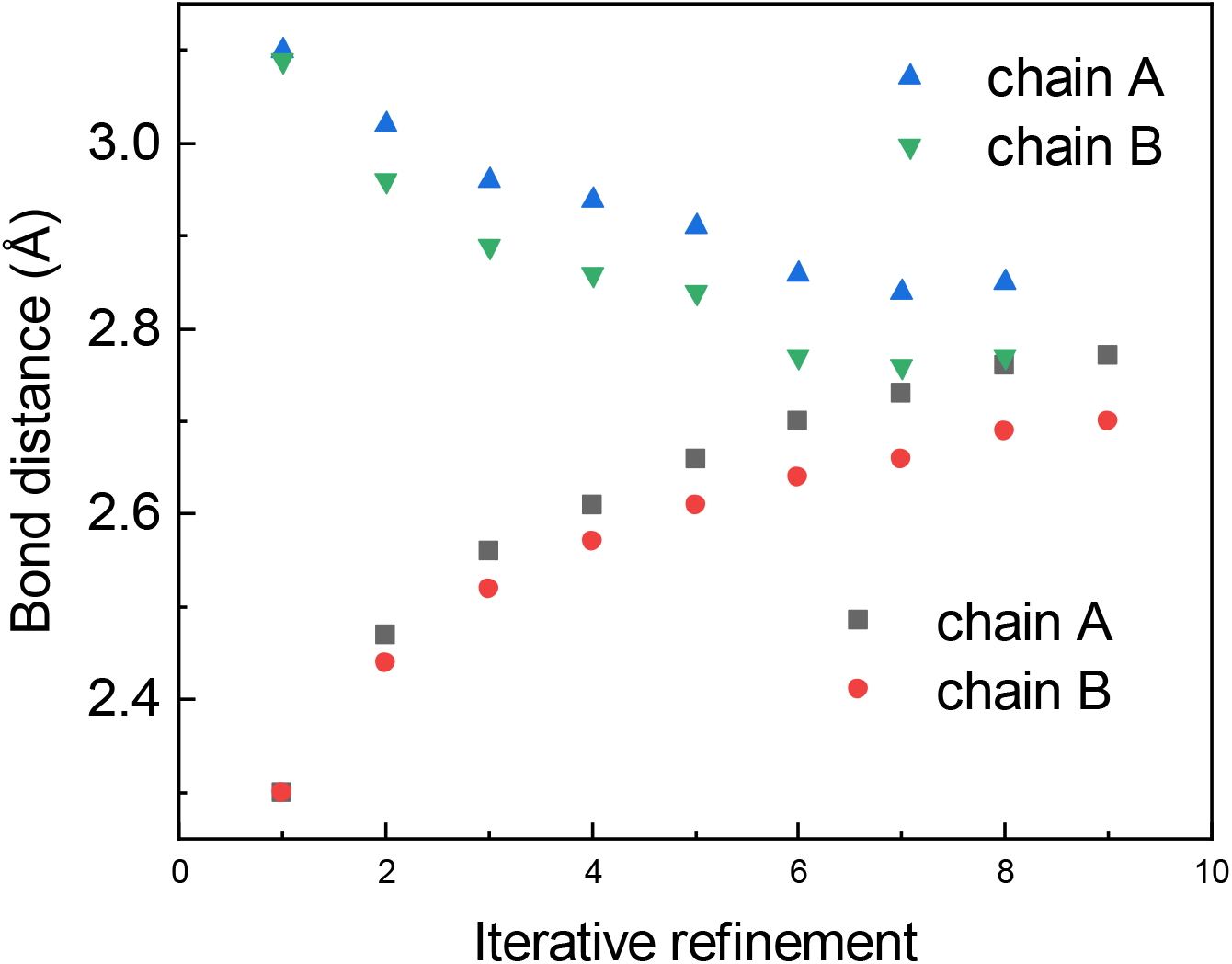
Bond distances between the Fe_d_ and thiol of C367 in interactive refinement cycles. The first cycles were performed with strict restraint in bond distances of either 2.3 Å or 3.1 Å between the Fe_d_ and thiol of C367 with standard deviation (σ) of 0.005 Å. In following interactive cycles, the refined model from cycle n was used as the input model in the n+1 cycle. Additionally, the refined distance between the Fe_d_ and thiol of C367 obtained in refinement cycle n was used as restraint in cycle n+1 with a very relaxed standard deviation (σ=0.1 Å).

DFT calculations were performed to complement the crystallographic analysis by assessing plausible H_inact_ structural models against the experimental geometry. Different ligands at the Fe_d_ were systematically explored: (i) deprotonated C367 (C367-S^−^), (ii) OH^−^ or (iii) H_2_O in the presence of protonated C367, and oxidized cysteine species, namely (iv) sulfenate (C367–SO^−^) and (v) sulfenic acid (C367–SOH) (Figure 3). Unlike the earlier study by Corrigan et al.,^*27*^ where the candidate ligands (including the C367 ligand, modeled as SCH_3_^−^) were treated as isolated molecular fragments, our calculations use an extended cluster model of the protein environment (Figure S4). Indeed, the higher-resolution Cb5AH structure available here enabled more realistic ligand placement and explicit evaluation of protein-matrix constraints around the H-cluster and C367. Selected geometric parameters for the optimized structures are reported in Figure 3 and Table S2, with particular focus on the Fe_d_–S(C367) distance and, for oxygen-based ligands, the Fe_d_–O(ligand) distance. Geometry optimizations were first performed with the Cα atom of C367 fixed at its crystallographic position and then repeated after releasing this constraint. Irrespective of the optimization protocol, only the C367–S^−^ model reproduced Fe_d_–S distances close to the experimental value of 2.77 Å. With the Cα atom constrained, the average Fe_d_–S distance was 2.58 Å across the tested functionals, whereas relaxation of the cysteine side chain shortened this distance to 2.45 Å. This trend is consistent with the tendency of the Fe–S bond to relax toward its typical equilibrium value in the absence of external constraints. However, such shortening may be likely an artifact arising from the limited size of the cluster model, which does not fully capture the restraining effect of the protein matrix. To test this hypothesis, a larger cluster model was constructed by including additional residues in proximity to C367 (Figures S4 and S5). In this extended model, optimizations performed without constraining the Cα atom of C367 yielded Fe_d_–S distances in the range of 2.60–2.62 Å (except for B3LYP-D4, which predicts a slightly shorter value of 2.551 Å), showing overall a better agreement with the experimental value than the smaller model (Figures 3 and S5). This supports the idea that the protein environment may help modulate the Fe–S interaction, weakening and making more reversible the coordination of C367 to Fe_d_. By contrast, oxygen-based ligands give geometries that deviate substantially from the experimental structure. Both the Fe_d_–O and Fe_d_–S distances differ markedly from the 2.67 Å reference value and show limited sensitivity to the applied constraints. For the H_2_O and OH^−^ ligand models, the SELF^*38*^ analysis (Steric Exclusion Localization Function, Figure S6) indicates steric strain between the Fe_d_-bound ligand and the C367 S–H group. Consistently, the optimized structures show a reorientation of the cysteine side chain, with the sulfur atom moving away from the ligand to reduce unfavorable contact. Upon release of the C367 Cα constraint, this relaxation becomes more pronounced: the Fe_d_–Cα(C367) distance slightly increases by ∼0.12–0.24 Å relative to the starting geometry (Figure S5), and the ligand shifts further away from the S–H group (Figure S6). Although these rearrangements partially relieve the steric strain, the ligand remains confined between the Fe_d_ and C367. This suggests that accommodation of a small oxygen-based ligand in this position is structurally unfavorable, further supporting the assignment of OH^−^ and H_2_O as less likely H_inact_ ligand candidates.

**Figure 3.**
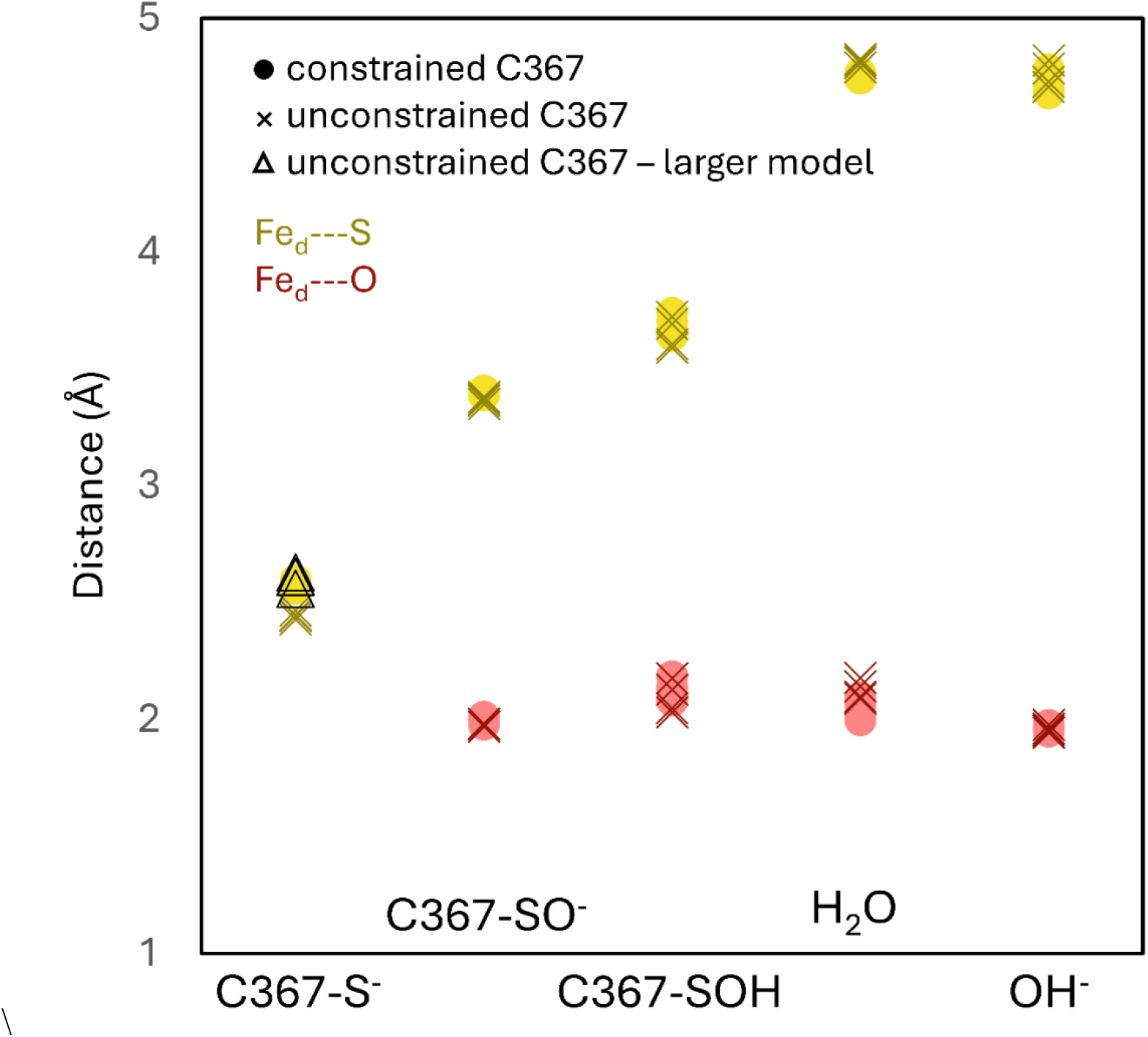
Selected geometrical parameters for the optimized H_inact_ candidate structures obtained with the four tested schemes: BP86-D4/def-TZVP, TPSS-D4/def-TZVP, TPSSh-D4/def-TZVP, and B3LYP-D4/def-TZVP. The Fe_d_–S(C367) distances are shown in yellow, whereas the Fe_d_–O distances are shown in red for models containing an oxygen-coordinating ligand. Distances are reported in Å. Values obtained with C367 constrained at the Cα atom are shown as dots, while those obtained with the C367 side chain fully relaxed are shown as crosses. For the C367–S^−^ model, the Fe_d_–S(C367) distances obtained with the larger cluster model are also reported and indicated by open triangles with a black outline. The data appear largely superimposed because the optimized distances are highly consistent across the tested functionals.

The analysis of the structure of CbA5H in the H_inact_ state with a resolution of 2.15 Å conclusively reveals that the ligand that binds to Fe_d_ is the thiol of C367, which is in line with our earlier proposal.^*28*^ This conclusion is well supported by new DFT calculations that employed the model of this CbA5H structure that we could refined to a higher resolution than previous structures.^*28*^ The distance of 2.77 Å bond between the Fe_d_ and the thiol that we obtained from systematic refinement is significantly longer than the typical length of 2.1-2.3 Å of a covalent bond between Fe and S in natural iron-sulfur clusters and synthetic iron-sulfur complexes.^*37, 39*^ A distance of 2.48 Å of the bond of the iron-axial-thiolate, -which is very similar to the examined case here-, in chloroperoxidase compound I was reported.^*40*^ When a thiolate is in a bridging position between two iron ions, the bond length is generally longer, namely about 2.44-2.51 Å. In a rare case, it can be up to 2.67 Å, as reported in a thiolate iron complex.^*41*^ Of important note is the rapid reversibility of the H_inact_ state of CbA5H,^*26-28*^ which requires a facile break of the bond once one-electron reduction occurs on [2Fe]_H,_ in contrast to stable covalent bonds. We hypothesize that this rapid reversibility of H_inact_ in CbA5H may be correlated with the exceptionally long bond distance between the Fe_d_ and the thiol of C367. A previous crystal structure of of DdH in the H_inact_ state reported a bond distance between the Fe_d_ and the extrinsic sulfide of about 2.4 Å, although the H_inact_ state of DdH appears to be as readily reversible as the H_inact_ of CbA5H. However, a lower occupancy (60 %) of the [2Fe]_H_ and the bound extrinsic sulfide as well as unexplained densities of the thiol group of C178 (corresponding to C367 of CbA5H) locating 2.79 Å to the extrinsic sulfide may have hindered an accurate determination of the bond distance.^*14*^ A structure of good resolution with high occupancies of the [2Fe]_H_ and the extrinsic sulfide would be more comparable with the structure of CbA5H refined here. Alternatively, as shown in our DFT calculations, the restraints from the protein matrix of C367 exerted a positive effect in “elongating” this Fe_d_-S bond distance, which may essentially explain the actual differences in the Fe_d_-S bond distances of CbA5H and DdH. A similar cysteine-thiolate binding to the Fe_d_ upon the formation of the H_inact_ state of ToHydA was proposed.^*32*^ However, compared to CbA5H, a significantly enhanced stability to O_2_ and also an enhanced stability of H_inact_ was observed in ToHydA.^*32*^ A shorter bond distance in H_inact_ of ToHydA may be one of the reasons that can explain the observations. Determining and comparing the precise geometries of the [2Fe]_H_ in O_2_-stable H_inact_ state from different [FeFe]-hydrogenases would help enhance understandings in O_2_ protection mechanism of [FeFe]-hydrogenases.

## ASSOCIATED CONTENT

The UniProtKB accession ID of CbA5H from *Clostridium beijerinckii* is A0A1I9RYV3.

## Supporting Information

The Supporting Information is available free of charge. Materials and methods, more structural modeling and DFT calculation details

## Author Contributions

J.D. and T.H. designed the project. J.D. and E.H. performed the structural analysis.

A.R. isolated and crystallized the protein. F.A. and C.G. conducted DFT calculations.

J.D. and F.A. drafted the manuscript. A.R., C.G. and T.H. edited the manuscript.

## Notes

The optimal resolution of 2.15 Å of the crystallographic data was determined using paired refinement^*42, 43*^(see details in the method). The structural results were obtained based on this 2.15 Å dataset if not stated otherwise. Using a more traditional cutoff method (I/σ ≥1 and CC1/2 ≥30%), the data were cut to 2.4 Å. Although the electron densities were weaker in the 2.4 Å dataset, all the structural results are still valid and very consistent with the results obtained from the 2.15 Å dataset. Results obtained from the 2.4 Å dataset corresponding to those in Figures 1 and 2 are shown in Figures S7 and S8. During revision of this study, we noticed some similar results were published by others.^47^

## ACKNOWLEDGMENT

J.D. thanks the German Research Foundation (Deutsche Forschungsgemeinschaft, DFG) with project number 461338801) for financial support. E.H. and T.H. received funding from the DFG Research Training Group GRK 2341 “Microbial Substrate Conversion (MiCon)”. The authors thank X-ray facility ESRF for data collection at beamline ID29.

## For Table of Contents use only

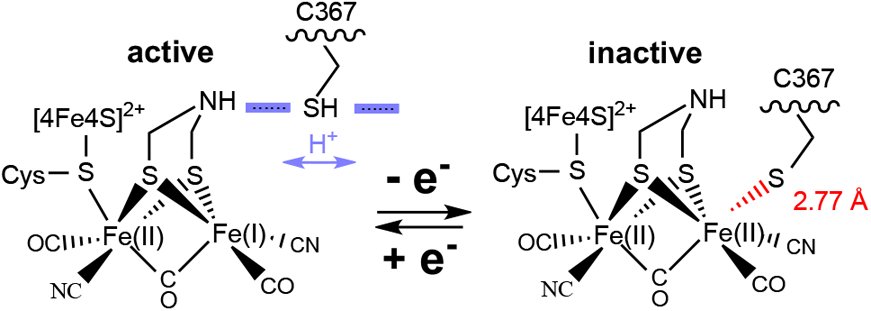

## REFERENCES

[1] Glick, B. R.; Martin, W. G.; and Martin, S. M. (1980) Purification and properties of the periplasmic hydrogenase from Desulfovibrio desulfuricans, Can J Microbiol 26, 1214–1223.

[2] Stripp, S. T.; Duffus, B. R.; Fourmond, V.; Léger, C.; Leimkühler, S.; Hirota, S.; Hu, Y.; Jasniewski, A.; Ogata, H.; and Ribbe, M. W. (2022) Second and outer coordination sphere effects in nitrogenase, hydrogenase, formate dehydrogenase, and CO dehydrogenase, Chemical reviews 122, 11900–11973.

[3] Peters, J. W.; Lanzilotta, W. N.; Lemon, B. J.; and Seefeldt, L. C. (1998) X-ray crystal structure of the Fe-only hydrogenase (CpI) from Clostridium pasteurianum to 1.8 angstrom resolution, Science 282, 1853–1858.

[4] Nicolet, Y.; Piras, C.; Legrand, P.; Hatchikian, C. E.; and Fontecilla-Camps, J. C. (1999) Desulfovibrio desulfuricans iron hydrogenase: the structure shows unusual coordination to an active site Fe binuclear center, Structure 7, 13–23.

[5] Pierik, A. J.; Hulstein, M.; Hagen, W. R.; and Albracht, S. P. (1998) A low-spin iron with CN and CO as intrinsic ligands forms the core of the active site in [Fe]-hydrogenases, The FEBS Journal 258, 572–578.

[6] Silakov, A.; Wenk, B.; Reijerse, E.; and Lubitz, W. (2009) 14N HYSCORE investigation of the H-cluster of [FeFe] hydrogenase: evidence for a nitrogen in the dithiol bridge, Physical Chemistry Chemical Physics 11, 6592–6599.

[7] Berggren, G.; Adamska, A.; Lambertz, C.; Simmons, T. R.; Esselborn, J.; Atta, M.; Gambarelli, S.; Mouesca, J. M.; Reijerse, E.; Lubitz, W.; Happe, T.; Artero, V.; and Fontecave, M. (2013) Biomimetic assembly and activation of [FeFe]-hydrogenases, Nature 499, 66–69.

[8] Esselborn, J.; Lambertz, C.; Adamska-Venkates, A.; Simmons, T.; Berggren, G.; Noth, J.; Siebel, J.; Hemschemeier, A.; Artero, V.; Reijerse, E.; Fontecave, M.; Lubitz, W.; and Happe, T. (2013) Spontaneous activation of [FeFe]-hydrogenases by an inorganic [2Fe] active site mimic, Nat. Chem. Biol. 9, 607–609.

[9] Mulder, D. W.; Guo, Y.; Ratzloff, M. W.; and King, P. W. (2017) Identification of a Catalytic Iron-Hydride at the H-Cluster of [FeFe]-Hydrogenase, J. Am. Chem. Soc. 139, 83–86.

[10] Reijerse, E. J.; Pham, C. C.; Pelmenschikov, V.; Gilbert-Wilson, R.; Adamska-Venkatesh, A.; Siebel, J. F.; Gee, L. B.; Yoda, Y.; Tamasaku, K.; and Lubitz, W. (2017) Direct Observation of an Iron-Bound Terminal Hydride in [FeFe]-Hydrogenase by Nuclear Resonance Vibrational Spectroscopy, J. Am. Chem. Soc. 139, 4306–4309.

[11] Winkler, M.; Senger, M.; Duan, J.; Esselborn, J.; Wittkamp, F.; Hofmann, E.; Apfel, U.-P.; Stripp, S. T.; and Happe, T. (2017) Accumulating the hydride state in the catalytic cycle of [FeFe]-hydrogenases, Nature communications 8, 16115.

[12] Lorent, C.; Katz, S.; Duan, J.; Kulka, C. J.; Caserta, G.; Teutloff, C.; Yadav, S.; Apfel, U.-P.; Winkler, M.; and Happe, T. (2020) Shedding light on proton and electron dynamics in [FeFe] hydrogenases, J. Am. Chem. Soc. 142, 5493–5497.

[13] Lemon, B. J.; and Peters, J. W. (2000) Photochemistry at the active site of the carbon monoxide inhibited form of the iron-only hydrogenase (CpI), J. Am. Chem. Soc. 122, 3793–3794.

[14] Rodríguez-Maciá, P.; Galle, L. M.; Bjornsson, R.; Lorent, C.; Zebger, I.; Yoda, Y.; Cramer, S. P.; DeBeer, S.; Span, I.; and Birrell, J. A. (2020) Caught in the Hinact: Crystal Structure and Spectroscopy Reveal a Sulfur Bound to the Active Site of an O2-stable State of [FeFe] Hydrogenase, Angewandte Chemie International Edition 59, 16786–16794.

[15] Duan, J.; Hemschemeier, A.; Burr, D. J.; Stripp, S. T.; Hofmann, E.; and Happe, T. (2023) Cyanide Binding to [FeFe]-Hydrogenase Stabilizes the Alternative Configuration of the Proton Transfer Pathway, Angewandte Chemie International Edition 62, e202216903.

[16] Martini, M. A.; Bikbaev, K.; Pang, Y.; Lorent, C.; Wiemann, C.; Breuer, N.; Zebger, I.; DeBeer, S.; Span, I.; and Bjornsson, R. (2023) Binding of exogenous cyanide reveals new active-site states in [FeFe] hydrogenases, Chemical Science 14, 2826–2838.

[17] Cornish, A. J.; Gartner, K.; Yang, H.; Peters, J. W.; and Hegg, E. L. (2011) Mechanism of proton transfer in [FeFe]-hydrogenase from Clostridium pasteurianum, J. Biol. Chem. 286, 38341–38347.

[18] Duan, J.; Senger, M.; Esselborn, J.; Engelbrecht, V.; Wittkamp, F.; Apfel, U. P.; Hofmann, E.; Stripp, S. T.; Happe, T.; and Winkler, M. (2018) Crystallographic and spectroscopic assignment of the proton transfer pathway in [FeFe]-hydrogenases, Nature communications 9, 4726.

[19] Senger, M.; Eichmann, V.; Laun, K.; Duan, J. F.; Wittkamp, F.; Knor, G.; Apfel, U. P.; Happe, T.; Winkler, M.; Heberle, J.; and Stripp, S. T. (2019) How [FeFe]-Hydrogenase Facilitates Bidirectional Proton Transfer, J. Am. Chem. Soc. 141, 17394–17403.

[20] Lampret, O.; Duan, J.; Hofmann, E.; Winkler, M.; Armstrong, F. A.; and Happe, T. (2020) The roles of long-range proton-coupled electron transfer in the directionality and efficiency of [FeFe]-hydrogenases, Proc Natl Acad Sci U S A 117, 20520–20529.

[21] Bruska, M. K.; Stiebritz, M. T.; and Reiher, M. (2011) Regioselectivity of H cluster oxidation, J. Am. Chem. Soc. 133, 20588–20603.

[22] Swanson, K. D.; Ratzloff, M. W.; Mulder, D. W.; Artz, J. H.; Ghose, S.; Hoffman, A.; White, S.; Zadvornyy, O. A.; Broderick, J. B.; and Bothner, B. (2015) [FeFe]-hydrogenase oxygen inactivation is initiated at the H cluster 2Fe subcluster, J. Am. Chem. Soc. 137, 1809–1816.

[23] Kubas, A.; Orain, C.; De Sancho, D.; Saujet, L.; Sensi, M.; Gauquelin, C.; Meynial-Salles, I.; Soucaille, P.; Bottin, H.; and Baffert, C. (2017) Mechanism of O2 diffusion and reduction in FeFe hydrogenases, Nature chemistry 9, 88–95.

[24] Mebs, S.; Kositzki, R.; Duan, J.; Kertess, L.; Senger, M.; Wittkamp, F.; Apfel, U.-P.; Happe, T.; Stripp, S. T.; and Winkler, M. (2018) Hydrogen and oxygen trapping at the H-cluster of [FeFe]-hydrogenase revealed by site-selective spectroscopy and QM/MM calculations, Biochimica et Biophysica Acta (BBA)-Bioenergetics 1859, 28–41.

[25] Esselborn, J.; Kertess, L.; Apfel, U.-P.; Hofmann, E.; and Happe, T. (2019) Loss of specific active-site iron atoms in oxygen-exposed [FeFe]-hydrogenase determined by detailed X-ray structure analyses, J. Am. Chem. Soc. 141, 17721–17728.

[26] Morra, S.; Arizzi, M.; Valetti, F.; and Gilardi, G. (2016) Oxygen stability in the new [FeFe]-hydrogenase from Clostridium beijerinckii SM10 (CbA5H), Biochemistry 55, 5897–5900.

[27] Corrigan, P. S.; Tirsch, J. L.; and Silakov, A. (2020) Investigation of the unusual ability of the [FeFe] hydrogenase from Clostridium beijerinckii to access an O2-protected state, J. Am. Chem. Soc. 142, 12409–12419.

[28] Winkler, M.; Duan, J.; Rutz, A.; Felbek, C.; Scholtysek, L.; Lampret, O.; Jaenecke, J.; Apfel, U.-P.; Gilardi, G.; and Valetti, F. (2021) A safety cap protects hydrogenase from oxygen attack, Nature communications 12, 756.

[29] Duan, J.; Rutz, A.; Kawamoto, A.; Naskar, S.; Edenharter, K.; Leimkuhler, S.; Hofmann, E.; Happe, T.; and Kurisu, G. (2025) Structural determinants of oxygen resistance and Zn2+-mediated stability of the [FeFe]-hydrogenase from Clostridium beijerinckii, Proc Natl Acad Sci U S A 122, e2416233122.

[30] Roseboom, W.; De Lacey, A. L.; Fernandez, V. M.; Hatchikian, E. C.; and Albracht, S. P. (2006) The active site of the [FeFe]-hydrogenase from Desulfovibrio desulfuricans. II. Redox properties, light sensitivity and CO-ligand exchange as observed by infrared spectroscopy, Journal of Biological Inorganic Chemistry 11, 102–118.

[31] Rodriguez-Macia, P.; Reijerse, E. J.; van Gastel, M.; DeBeer, S.; Lubitz, W.; Rudiger, O.; and Birrell, J. A. (2018) Sulfide Protects [FeFe] Hydrogenases From O2, J. Am. Chem. Soc. 140, 9346–9350.

[32] Ghosh, S.; Das, C. K.; Uddin, S.; Stripp, S. T.; Engelbrecht, V.; Winkler, M.; Leimkuhler, S.; Brocks, C.; Duan, J.; Schafer, L. V.; and Happe, T. (2025) Protein Dynamics Affect O2-Stability of Group B [FeFe]-Hydrogenase from Thermosediminibacter oceani, J. Am. Chem. Soc. 147, 15170–15180.

[33] Fasano, A.; Jacq-Bailly, A.; Wozniak, J.; Fourmond, V.; and Léger, C. (2024) Catalytic bias and redox-driven inactivation of the group B FeFe hydrogenase CpIII, ACS Catalysis 14, 7001–7010.

[34] Kisgeropoulos, E. C.; Ratzloff, M. W.; Stroeva-Dahl, E. M.; Hasan, S.; Varghese, F.; Artz, J. H.; Peters, J. W.; Mulder, D. W.; and King, P. W. (2025) H-cluster Intermediates and Catalytic Properties of Clostridium pasteurianum [FeFe]-Hydrogenase III, Biochemistry 64, 2455–2466.

[35] Felbek, C.; Arrigoni, F.; de Sancho, D.; Jacq-Bailly, A.; Best, R. B.; Fourmond, V.; Bertini, L.; and Léger, C. (2021) Mechanism of hydrogen sulfide-dependent inhibition of FeFe hydrogenase, ACS Catalysis 11, 15162–15176.

[36] Corrigan, P. S.; Majer, S. H.; and Silakov, A. (2023) Evidence of atypical structural flexibility of the active site surrounding of an [FeFe] hydrogenase from Clostridium beijerinkii, J. Am. Chem. Soc. 145, 11033–11044.

[37] Krest, C. M.; Silakov, A.; Rittle, J.; Yosca, T. H.; Onderko, E. L.; Calixto, J. C.; and Green, M. T. (2015) Significantly shorter Fe–S bond in cytochrome P450-I is consistent with greater reactivity relative to chloroperoxidase, Nature chemistry 7, 696–702.

[38] Just, G. H.; Lefebvre, C.; Rajamani, A.; Khartabil, H.; Pilmé, J.; and Hénon, É. (2026) A visualizable and widely applicable steric repulsion descriptor for guiding experimental chemistry, Chemical Science.

[39] Venkateswara Rao, P.; and Holm, R. (2004) Synthetic analogues of the active sites of iron-sulfur proteins, Chemical reviews 104, 527–560.

[40] Stone, K. L.; Behan, R. K.; and Green, M. T. (2005) X-ray absorption spectroscopy of chloroperoxidase compound I: Insight into the reactive intermediate of P450 chemistry, Proceedings of the National Academy of Sciences 102, 16563–16565.

[41] Krishnamurthy, D.; Sarjeant, A. N.; Goldberg, D. P.; Caneschi, A.; Totti, F.; Zakharov, L. N.; and Rheingold, A. L. (2005) Mononuclear, dinuclear, and pentanuclear [{N, S (thiolate)} Iron (II)] complexes: nuclearity control, incorporation of hydroxide bridging ligands, and magnetic behavior, Chemistry–A European Journal 11, 7328–7341.

[42] Maly, M.; Diederichs, K.; Dohnalek, J.; and Kolenko, P. (2020) Paired refinement under the control of PAIREF, IUCrJ 7, 681–692.

[43] Maly, M.; Diederichs, K.; Dohnalek, J.; and Kolenko, P. (2021) PAIREF: paired refinement also for Phenix users, Acta Crystallogr F Struct Biol Commun 77, 226–229.

[44] Alogaidi, A.; Carr, S. B.; Hudson, L.; Lloyd-Laney H.; Parkin A.; Love, A.; George, M. W.; Pordea A.; Morra, Simone. Probing the role of accessory domains in oxygen stability of [FeFe]-hydrogenases. J. Am. Chem. Soc. 2026 doi.org/10.1021/jacs.6c05865

