## supporting information for "Direct Binding of Cysteine-367 Thiolate to the Active Site of the [FeFe]-Hydrogenase from *Clostridium beijerinckii* in the O_2_-stable State"

*Corresponding author:

### Content


Materials, methods and notes S3

Figure S1. Electron density maps of the H-cluster and C367 of CbA5H in the $H_{inact}$ state in chain B. S6

Figure S2. Modeling a hydroxide ion between the $Fe_d$ and the thiol of C367. S7

Figure S3. Comparison of density maps of the $[2Fe]_H$ and C367 when modeling different distances between the $Fe_d$ and the thiol of C367. S8

Figure S4. CbA5H models used for DFT geometry optimizations. S9

Figure S5. DFT optimized structures of $H_{inact}$ candidates. S10

Figure S6. SELF analysis of interfragment steric overlap between the C367 S–H group and the $Fe_d$-bound ligand in $OH^-$ and $H_2O$ models. S11

Figure S7. Electron density maps of the H-cluster and C367 of CbA5H in the $H_{inact}$ state in chain A based on the 2.4 Å dataset. S13

Figure S8. Bond distances between the Fed and thiol of C367 in interactive refinement cycles based on the 2.4 Å dataset. S14

Table S1 X-ray data collection and refinement statistics S15

Table S2. Optimized $Fe_d$-ligand distances. S16

Table S3. Natural population analysis (NPA) of $H_{inact}$. S17

References S18

### Materials, methods and notes

The raw diffraction data of 6TTL[1] were reprocessed using XDS[2] package. The extension of the resolution from previous 2.9 Å to current 2.15 Å can be explained by two main reasons. First, the new version of XDS released in recent years allows for better distinguishing signals of weak intensities at high resolutions from noise. Applying the same input parameters, processing with the newer version resulted in the resolution improvement from 2.9 Å to 2.35 Å. Second, the parameter of "BACKGROUND_RANGE" was set to use initial two hundred instead of four images that was a default setting and used in our previous data processing. After combing both, the data were processed to 2.08 Å with fulfilling CC1/2 > 0.1 and I/σ > 1. An optimal resolution of 2.15 Å was determined by performing paired refinement using PAIREF.[3, 4] The procedure is briefly outlined as follows. The processed diffraction data were cut to 2.9 Å and stepwise (in a step of 0.2 Å, i.e. 2.7 Å, 2.5 Å, 2.3 Å and 2.1 Å) added the high-resolution data for refinement using PDB 9F47[5] as the starting input model. The results showed that overall Rfree decreased until the data included up to 2.1 Å. Another cycle of paired refinement was performed with the starting resolution of 2.21 Å and a finer step of 0.03 Å. The results showed that overall Rfree decreased until the data included up to 2.15 Å, which is the final resolution. Interactive refinements in Phenix[6] were performed to optimize the model. Refined models were inspected and manually corrected using Coot.[7] Strict geometry restraints of the $[2Fe]_H$ derived from high resolution structures of CpI ([FeFe]-hydrogenase I from *Clostridium pasteurianum*)[8, 9] were used through the refinement. TLS (translation, libration and screw) refinement was applied. Automatic search in Phenix refinement was used to define the TLS groups. The groups were identified by Phenix.[6] Residues 1-113, 114-246 and 247-639 were defined as three different groups, which represent three different substructures, namely SLBB-, $Zn^{2+}$-binding and fused ferredoxin and H-cluster domains. The TLS group definitions are the same in chains A and B. The inclusion of TLS refinement resulted in drops of about 1% for both Rfree and Rwork. NCS (non-crystallographic symmetry) torsion-angle restraints were used during refinement.

### DFT calculations

Density functional theory (DFT) calculations were performed using TURBOMOLE 7.4.1.[10] The initial model was derived from the refined crystallographic structure at 2.08 Å resolution (chain B). A cluster model comprising 163 atoms was constructed by selecting residues to account for: (i) hydrogen-bonding interactions involving the Fe ligands, (ii) the steric shape of the binding pocket surrounding the distal iron site, and (iii) the local environment of C367, including neighboring residues to reproduce its structural constraints. All calculations employed the def-TZVP basis set.[11] Geometry optimizations were carried out using four exchange–correlation functionals (BP86[12, 13], B3LYP[14, 15], TPSS[16], and TPSSh[17]), including D4 dispersion corrections[18]. Protein/environment effects were treated using the COSMO model with a dielectric constant ε = 4.[19] During geometry optimizations, strategically selected atoms of the protein backbone were kept fixed to avoid unrealistic displacements (Figure S4). C367 was treated using two alternative schemes: (i) full relaxation within the constraints imposed by the surrounding residues included in the cluster, or (ii) partial constraint with the Cα atom fixed. The $H_{inact}$ state was modeled with an overall singlet spin state S = 0. The electronic structure of the $[4Fe4S]_H$ cubane subcluster was described using a broken-symmetry (BS) approach, consistent with magnetic coupling schemes adopted in previous studies.[20, 21] To further assess the effect of structural constraints, a larger cluster model (228 atoms) was also constructed (Figure S4 and S5). providing a more extensive representation of the protein environment around the active site and C367. Given the high consistency observed among $Fe_d$-ligand distances, the optimization of this model was limited to BP86.

Steric crowding in the $OH^-$ and $H_2O$ $H_{inact}$ models was analyzed using the Steric Exclusion Localization Function (SELF)[22] as implemented in IGMPlot[23]. This approach allows the identification of regions where short-range steric effects arise directly from the quantum-mechanical electron density. As representative cases, SELF analyses were performed on the electronic structures obtained with the BP86 functional for the $OH^-$ and $H_2O$ models, considering both optimizations with C367 constrained at the Cα atom and unconstrained (Figure S6). For each model, two fragments were defined: the S–H group of C367 and the exogenous ligand bound to $Fe_d$ ($OH^-$ or $H_2O$). The analysis was therefore focused on the interfragment contact between the protonated cysteine and the oxygen-based ligand located at the distal coordination site. The graphical representation was obtained by plotting an isosurface of the δginter/ρ descriptor, which identifies regions of electron density overlap between fragments, and mapping the corresponding SELF(r) values onto this surface. In this representation, the isosurface locates the interfragment overlap region, while the color encodes the local magnitude of the interfragment Pauli kinetic energy excess (SELF), which reflects the degree of electron density overlap and the associated steric (Pauli) repulsion. A δginter/ρ isovalue of 0.6 a.u.$^{-1}$ was used for both models. The SELF(r) color scale was kept identical in the two cases, ranging from 0 to 8 kcal mol$^{-1}$ bohr$^{-3}$, to ensure direct comparability. All visualization parameters were kept constant so that differences in the maps reflect differences in the computed interfragment steric interaction rather than graphical artifacts. The overall steric crowding between the selected fragments was quantified using the integrated SELF score, iSELF, obtained by numerical integration of the local SELF(r) function over the grid region considered in the analysis. Thus, while SELF(r) describes the local steric-repulsion density, reported in kcal mol$^{-1}$ bohr$^{-3}$, iSELF provides a global SELF-based steric-crowding score for the chosen fragment pair, reported in kcal $mol^{-1}$. The integrated iSELF values should be interpreted as SELF-based steric-crowding scores, obtained by integrating the local interfragment Pauli-repulsion density over the grid, rather than as conventional interaction energies.

**Figure S1. Electron density maps of the H-cluster and C367 of CbA5H in the H$_{inact}$ state in chain B.**

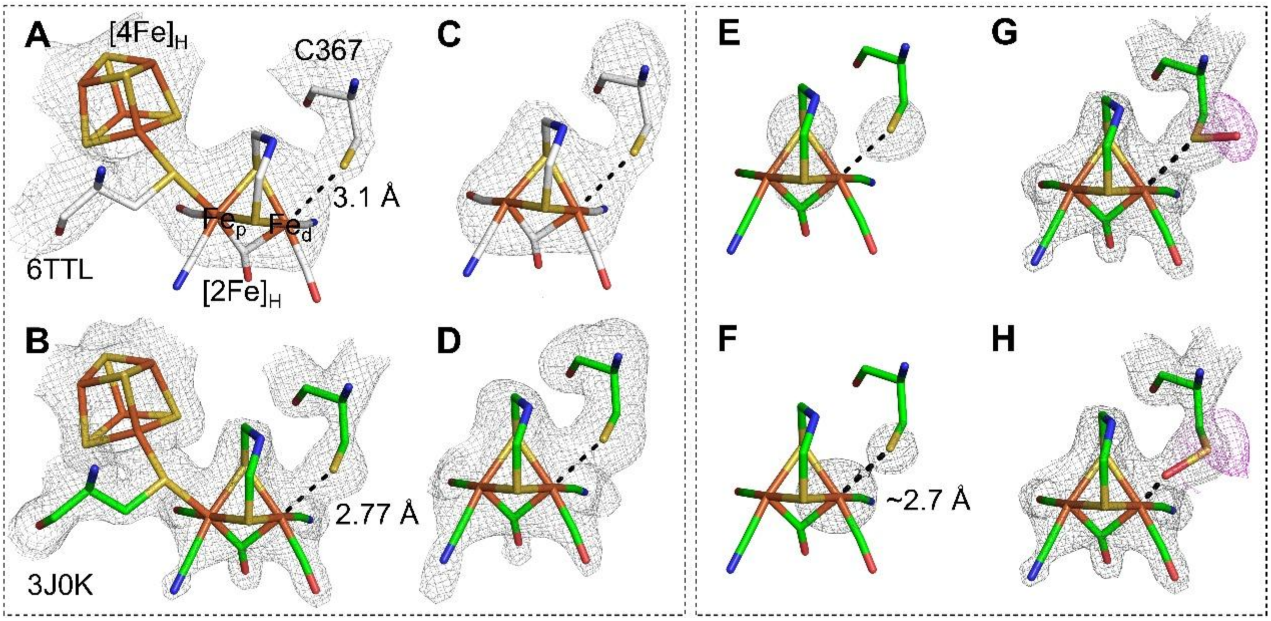


**Figure S1. Electron density maps of the H-cluster and C367 of CbA5H in the H$_{inact}$ state in chain B. A** and **B** show 2mFo-Fc maps of the H-cluster and C367 of CbA5H in the H$_{inact}$ state from structures of PDB 6TTL and the new structure presented here (PDB 30JK). Panels **A-D** show comparisons of density maps of the previous structure (6TLL: **A**/**C**) and the new structure (30JK) from this study with a focus on the H-cluster and C367. The two structures are distinguished by coloring carbon atoms white and green for 6TTL (**A**, **C**) and 30JK (**B**, **D**, **E-H**), respectively. **E** shows densities of three sulfur atoms, namely two from the dithiolate of the [2Fe]$_H$ and one from the thiol of C367. **F** shows individually calculated omitting maps of the Fe$_d$ of the [2Fe]$_H$ and thiol of C367. **G-H** show two modeling schemes of sulfenic acid at position 367. 2mFo-DFc maps and mFo-DFc omitting maps were colored gray and shown in **A**/**B**/**G**/**H** and **C**/**D**/**G**/**H**, respectively. Additional negative mFo-DFc maps colored magenta are shown in **G** and **H**. The contour levels of the maps are 2.5 $\sigma$ in **A**/**B**, 3.2 $\sigma$ in **C**, 4.5 $\sigma$ in **D**, 7 $\sigma$ in **E**, 9 $\sigma$ in **F** and 2.5 $\sigma$ in **G**/**H**. The distances between the Fe$_d$ and the thiol of C367 are indicated in **A**/**B**/**F**. The figure is prepared based on structural information from chain B.

**Figure S2. Modeling a hydroxide ion between the $Fe_d$ and the thiol of C367.**

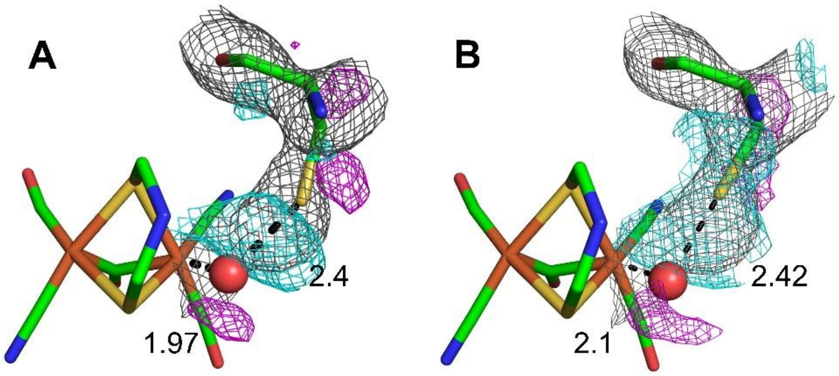


**Figure S2. Modeling a hydroxide ion between the $Fe_d$ and the thiol of C367.** The resulting 2mFo-DFc and mFo-DFc maps from chains A and B are shown in panels **A** and **B**, respectively, when a hydroxide ion shown as a red sphere was modeled between the $Fe_d$ and the thiol of C367. 2mFo-DFc maps colored gray are shown for the modeled hydroxide ions and C367. Positive and negative mFo-DFc maps of the modelled hydroxide ions and C367 were colored cyan and magenta, respectively. All the maps were contoured at 2.5 σ. The numbers indicate distances in Å between the modeled hydroxide and the $Fe_d$ or the thiol of C367.

**Figure S3. Comparison of density maps of the $[2Fe]_H$ and C367 when modeling different distances between the $Fe_d$ and the thiol of C367.**

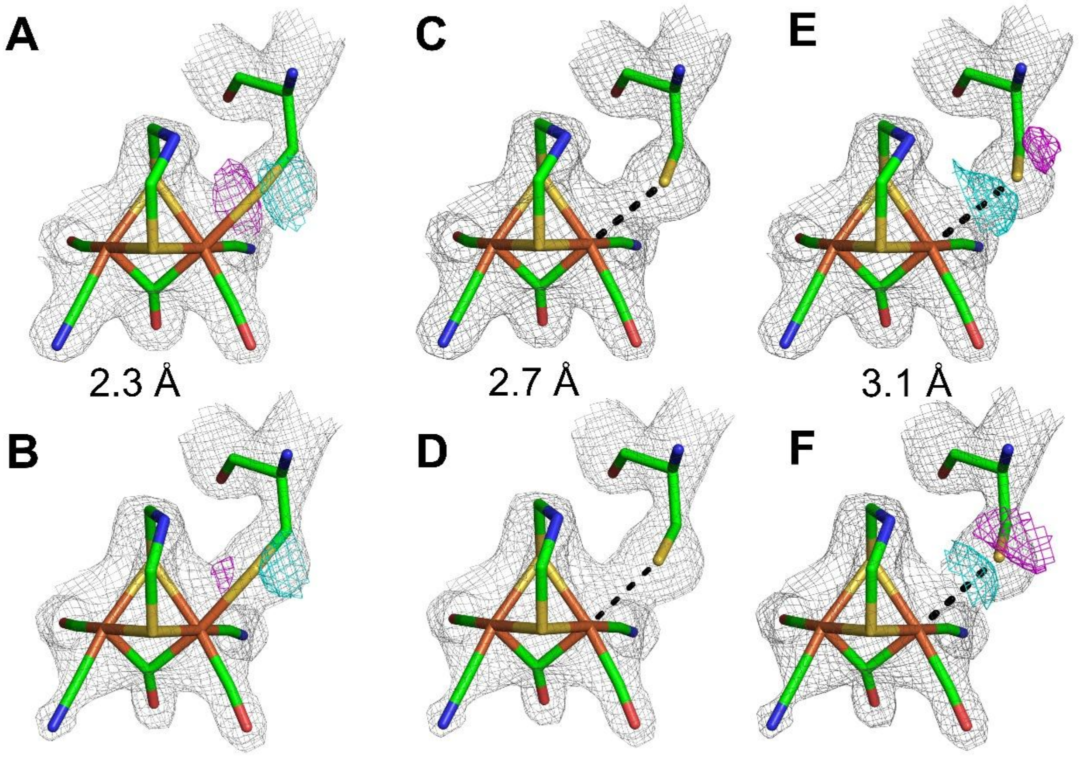


**Figure S3. Comparison of density maps of the $[2Fe]_H$ and C367 when modeling different distances between the $Fe_d$ and the thiol of C367.** **A**-**B**, **C**-**D** and **E**-**F** show the modeling results with restrained bond distances between the $Fe_d$ and thiol of C367 of 2.3 Å, 2.7 Å and 3.1 Å, respectively. **A**/**C**/**E** and **B**/**D**/**F** show results from chains A and B, respectively. 2mFo-DFc maps were colored gray in each panel. Positive and negative mFo-DFc maps were colored cyan and magenta respectively. All maps were contoured at 2.5 σ.

**Figure S4. CbA5H models used for DFT geometry optimizations.**

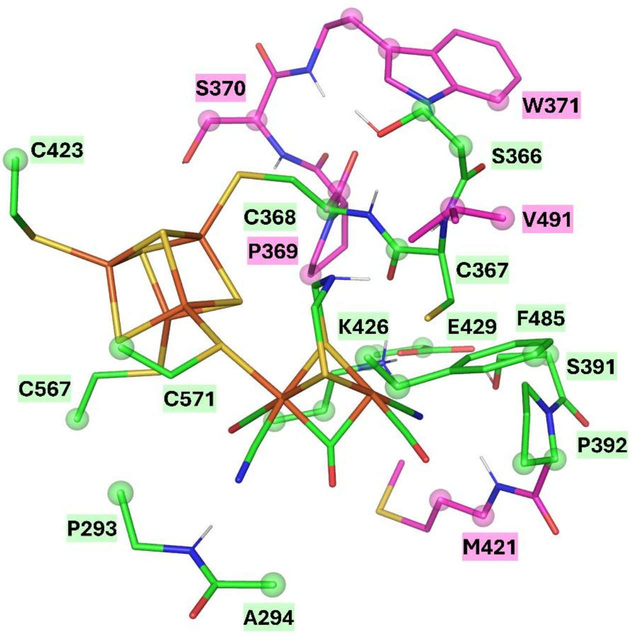


**Figure S4. CbA5H models used for DFT geometry optimizations.** The 163-atoms model is reported in green, while additional residues in the 228-atoms model are reported in pink. Atoms strategically constrained during optimizations are indicated with spheres. C367 has been treated using two alternative schemes: (i) full relaxation within the constraints imposed by the surrounding residues included in the cluster, or (ii) partial constraint with the Cα atom fixed.

**Figure S5. DFT optimized structures of $H_{inact}$ candidates.**

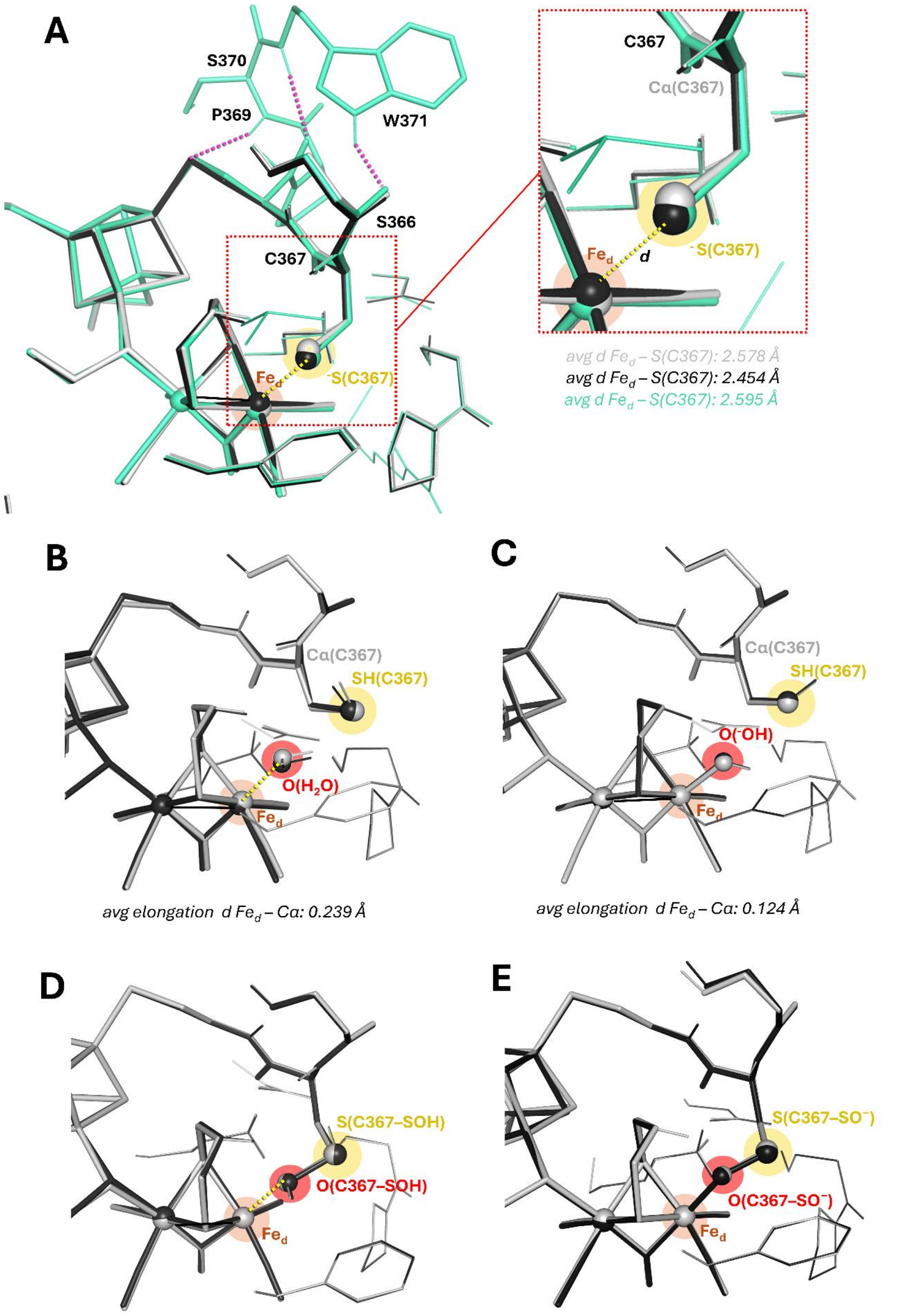


**Figure S5. DFT optimized structures of $H_{inact}$ candidates.** BP86-D4 optimized structures of the Hinact candidates, showing the $Fe_d$–ligand interaction: **A** C367–$S^-$, **B** $H_2O$, **C** $OH^-$, **D** sulfenic acid, and **E** sulfenate. Grey and black structures correspond to optimizations with C367 Cα constrained and fully relaxed, respectively. For the C367–$S^-$ model, the extended-cluster optimized structure is shown in cyan. $Fe_d$–S(C367) distances are reported together with the average values obtained across the four tested functionals for the smaller model. In the extended cluster model, the larger $Fe_d$–S(C367) distance of ~2.6 Å obtained after unconstrained optimization is likely due to the additional support provided by the 369–371 region, whose H-bonding network restrains the C367 environment and limits its relaxation toward $Fe_d$ (**A**).

**Figure S6. SELF analysis of interfragment steric overlap between the C367 S–H group and the $Fe_d$-bound ligand in $OH^-$ and $H_2O$ models.**

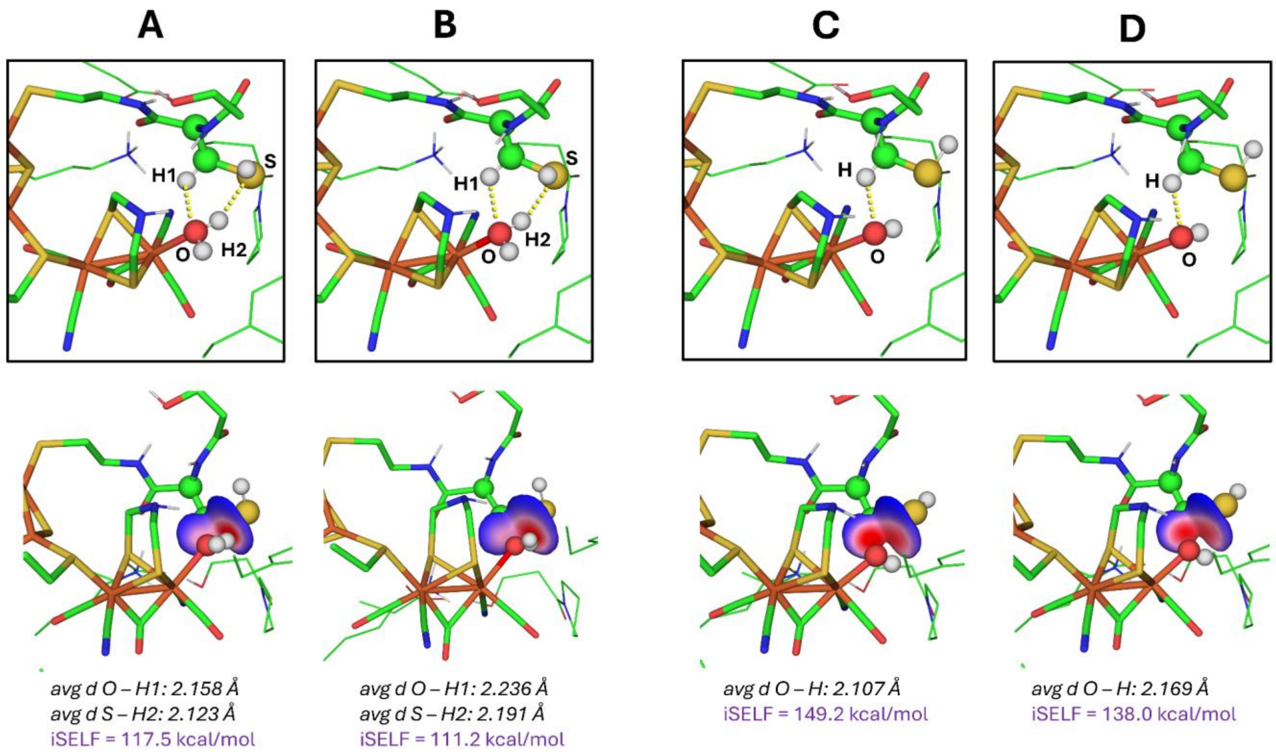


**Figure S6. SELF analysis of interfragment steric overlap between the C367 S–H group and the $Fe_d$-bound ligand in $OH^-$ and $H_2O$ models.** Panels **A** and **B** correspond to the $H_2O$ model, while panels C and D correspond to the $OH^-$ model. In panels **A** and **C**, the Cα atom of C367 is constrained, whereas in panels **B** and **D** the constraint is released, allowing structural relaxation. For each panel, the top image reports selected geometric parameters, while the bottom image shows the Steric Exclusion Localization Function (SELF) mapped on the $\delta g^{inter}/\rho$ isosurface. The $\delta g^{inter}/\rho = 0.6$ a.u.$^{-1}$ isosurfaces identify regions of interfragment electron density overlap between the C367 S–H group and the $Fe_d$-bound ligand. The surfaces are colored according to local SELF(r) values in the range 0–8 kcal mol$^{-1}$ bohr$^{-3}$, using the same isovalue and color scale for all panels to allow direct comparison. The color scale ranges from blue to pink/red, with red indicating regions of highest interfragment electron density overlap associated with strong Pauli (steric) repulsion. In optimized geometries of these models, the C367 sulfur atom rotates to adopt a geometry that maximizes its distance from the ligand, thereby reducing steric crowding. Upon relaxation of the C367 constraint (panels **B** and **D**), a rearrangement of both the ligand and the cysteine side chain is observed. The ligand shifts away from the S–H group. This is reflected in a decrease of the integrated iSELF values, from

117.5 to 111.2 kcal mol$^{-1}$ for the $H_2O$ model and from 149.2 to 138.0 kcal mol$^{-1}$ for the $OH^-$ model, indicating partial relief of interfragment steric strain.

**Figure S7. Electron density maps of the H-cluster and C367 of CbA5H in the H$_{inact}$ state in chain A based on the 2.4 Å dataset.**

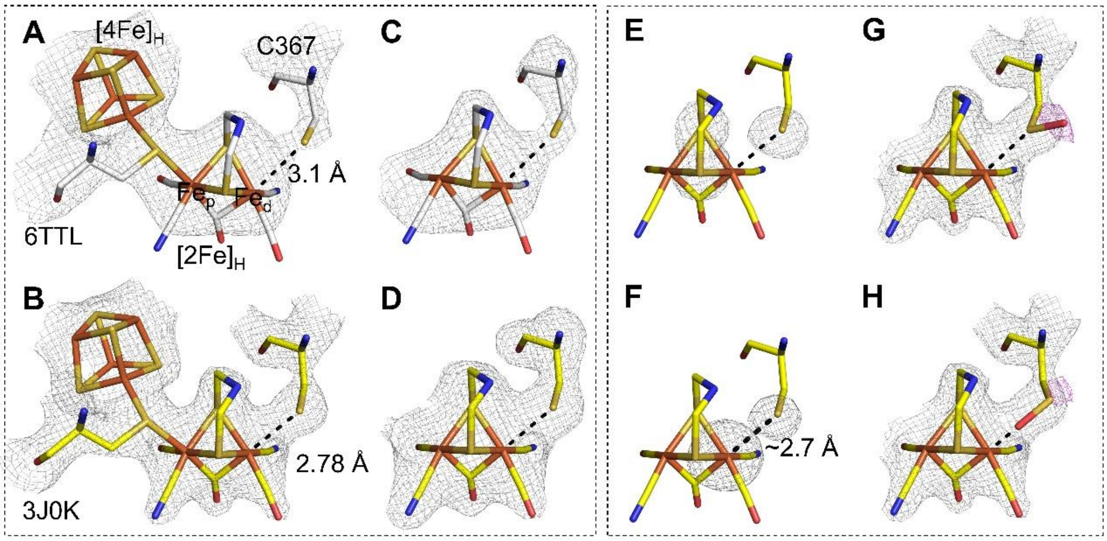


**Figure S7. Electron density maps of the H-cluster and C367 of CbA5H in the H$_{inact}$ state in chain A based on the 2.4 Å dataset.** **A** and **B** show 2mFo-Fc maps of the H-cluster and C367 of CbA5H in the H$_{inact}$ state from structures of PDB 6TTL and the new structure cut at 2.4 Å. Panels **A-D** show comparisons of density maps of the previous structure (6TLL: **A**/**C**) and the new structure (30JK) from this study with a focus on the H-cluster and C367. The two structures are distinguished by coloring carbon atoms white and yellow for 6TTL (**A**, **C**) and the 2.4 Å structure (**B**, **D**, **E-H**), respectively. **E** shows densities of three sulfur atoms, namely two from the dithiolate of the [2Fe]$_H$ and one from the thiol of C367. **F** shows individually calculated omitting maps of the Fe$_d$ of the [2Fe]$_H$ and thiol of C367. **G-H** show two modeling schemes of sulfenic acid at position 367. 2mFo-DFc maps and mFo-DFc omitting maps were colored gray and shown in **A**/**B**/**G**/**H** and **C**/**D**/**G**/**H**, respectively. Additional negative mFo-DFc maps colored magenta are shown in **G** and **H**. The contour levels of the maps are 2.5 $\sigma$ in **A**/**B**, 3.2 $\sigma$ in **C**, 4.5 $\sigma$ in **D**, 7 $\sigma$ in **E**, 9 $\sigma$ in **F** and 2.5 $\sigma$ in **G**/**H**. The distances between the Fe$_d$ and the thiol of C367 are indicated in **A**/**B**/**F**. The figure is prepared based on structural information from chain A.

**Figure S8. Bond distances between the Fed and thiol of C367 in interactive refinement cycles based on the 2.4 Å dataset.**

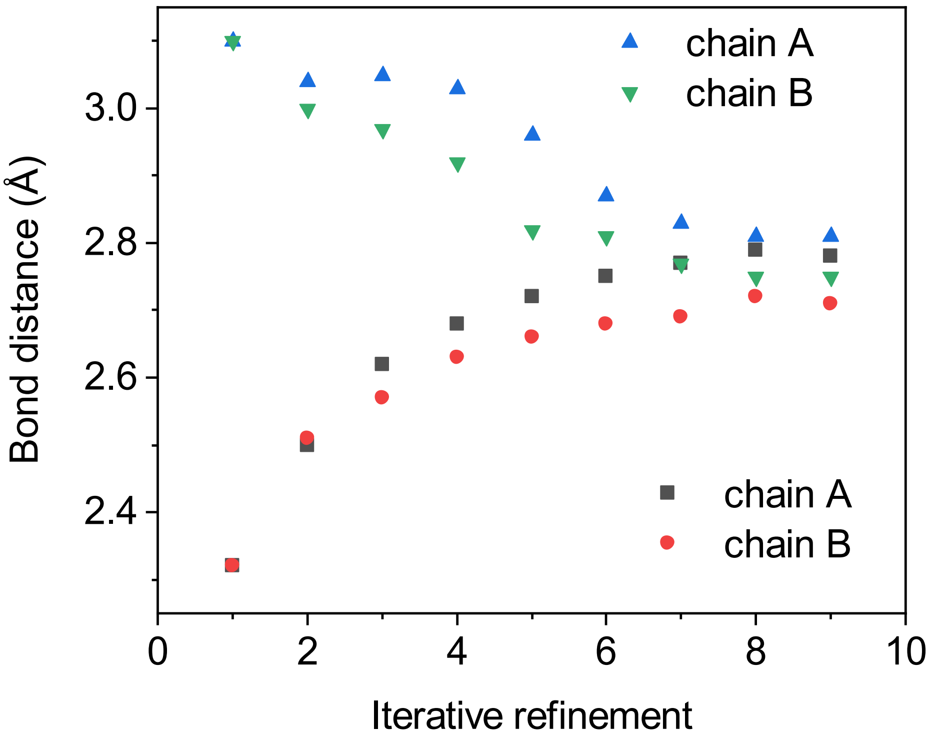


**Figure S8. Bond distances between the Fed and thiol of C367 in interactive refinement cycles based on the 2.4 Å dataset.** Bond distances between the Fed and thiol of C367 in interactive refinement cycles. The first cycles were performed with strict restraint in bond distances of either 2.3 Å or 3.1 Å between the Fed and thiol of C367 with standard deviation (σ) of 0.005 Å. In following interactive cycles, the refined model from cycle n was used as the input model in the n+1 cycle. Additionally, the refined distance between the Fed and thiol of C367 obtained in refinement cycle n was used as restraint in cycle n+1 with a very relaxed standard deviation (σ=0.1 Å). The averaged bond distance is 2.78±0.03 Å.

**Table S1 X-ray data collection and refinement statistics**

| | CbA5H | CbA5H |
|---|---|---|
| **PDB** | 30JK | |
| **Data collection** | | |
| X-ray source | ESRF-ID29 | ESRF-ID29 |
| Wavelength (Å) | 0.97625 | 0.97625 |
| Space group | $P\ 4_2\ 2_1\ 2$ | $P\ 4_2\ 2_1\ 2$ |
| Cell dimensions | | |
| a, b, c (Å) | 168.42, 168.42, 126.31 | 168.42, 168.42, 126.31 |
| α, β, γ (°) | 90.00, 90.00, 90.00 | 90.00, 90.00, 90.00 |
| Resolution (Å) | 49.07-2.15 (2.21-2.15)* | 49.07-2.4 (2.46-2.4)* |
| $R_{meas}$ | 0.464(4.68)* | 0.405(4.09)* |
| I / σ(I) | 8.97 (1.01)* | 11.88 (1.47)* |
| Completeness (%) | 98.8 (92.9)* | 99.4 (100)* |
| Redundancy | 42.6 (15.6)* | 50.8 (33.4)* |
| CC1/2 | 0.999 (0.131)* | 0.999 (0.3)* |
| **Refinement** | | |
| Resolution (Å) | 49.07-2.15 | |
| No. reflections | 97318 | |
| $R_{work}$ / $R_{free}$ | 0.2493/0.2796 | |
| No. atoms | | |
| Protein | 9463 | |
| Ligand | 84 | |
| Water/ions | 613/11 | |
| B-factors | | |
| Protein | 52.16 | |
| Ligand | 41.36 | |
| Water | 49.52 | |
| R.m.s deviations | | |
| Bond lengths (Å) | 0.003 | |
| Bond angles (°) | 0.592 | |
| Ramachandran plot | | |
| favored | 97.12 | |
| allowed | 2.88 | |
| outliers | 0 | |
| Clash score | 5.04 | |

*Numbers in brackets indicate values in the highest resolution shell

### Table S2. Optimized $Fe_d$-ligand distances.

| $H_{inact}$ model/ligand identity | Distances (Å) | *BP86-D4 def-TZVP COSMO ε=4* | *TPSS-D4 def-TZVP COSMO ε=4* | *TPSSh-D4 def-TZVP COSMO ε=4* | *B3LYP-D4 def-TZVP COSMO ε=4* |
|---|---|---|---|---|---|
| **A** | | | | | |
| **C367-$S^-$ (Cα constrained)** | **$Fe_d$-S(C367-$S^-$)** | 2.586 | 2.567 | 2.561 | 2.596 |
| **C367-$S^-$ (Cα relaxed)** | **$Fe_d$-S(C367-$S^-$)** | 2.476 | 2.443 | 2.430 | 2.467 |
| **C367-$S^-$ (larger model, Cα relaxed)** | **$Fe_d$-S(C367-$S^-$)** | 2.622 | 2.604 | 2.602 | 2.551 |
| **B** | | | | | |
| **$H_2O$ (Cα constrained)** | **$Fe_d$-S(C367-SH)** | 4.767 | 4.760 | 4.742 | 4.761 |
| | **$Fe_d$-O($H_2O$)** | 2.178 | 2.143 | 2.091 | 2.099 |
| **$H_2O$ (Cα relaxed)** | **$Fe_d$-S(C367-SH)** | 4.819 | 4.825 | 4.789 | 4.809 |
| | **$Fe_d$-O($H_2O$)** | 2.095 | 2.067 | 1.997 | 2.021 |
| **C** | | | | | |
| **$OH^-$ (Cα constrained)** | **$Fe_d$-S(C367-SH)** | 4.669 | 4.678 | 4.685 | 4.781 |
| | **$Fe_d$-O($OH^-$)** | 1.978 | 1.962 | 1.944 | 1.953 |
| **$OH^-$ (Cα relaxed)** | **$Fe_d$-S(C367-SH)** | 4.728 | 4.706 | 4.785 | 4.819 |
| | **$Fe_d$-O($OH^-$)** | 1.978 | 1.963 | 1.949 | 1.954 |
| **D** | | | | | |
| **C367-SOH (Cα constrained)** | **$Fe_d$-S(C367-SOH)** | 3.740 | 3.703 | 3.664 | 3.643 |
| | **$Fe_d$-O(C367-SOH)** | 2.186 | 2.136 | 2.081 | 2.087 |
| **C367-SOH (Cα relaxed)** | **$Fe_d$-S(C367-SOH)** | 3.723 | 3.693 | 3.590 | 3.605 |
| | **$Fe_d$-O(C367-SOH)** | 2.179 | 2.129 | 2.032 | 2.049 |
| **E** | | | | | |
| **C367-$SO^-$ (Cα constrained)** | **$Fe_d$-S(C367-$SO^-$)** | 3.405 | 3.391 | 3.387 | 3.408 |
| | **$Fe_d$-O(C367-$SO^-$)** | 2.013 | 1.992 | 1.979 | 1.992 |
| **C367-$SO^-$ (Cα relaxed)** | **$Fe_d$-S(C367-$SO^-$)** | 3.380 | 3.364 | 3.350 | 3.370 |
| | **$Fe_d$-O(C367-$SO^-$)** | 2.001 | 1.983 | 1.970 | 1.982 |

Optimized geometric parameters collected in Figure 3. Labels A–E refer to the panels shown in Figure S5.

**Table S3. Natural population analysis (NPA) of $H_{inact}$.**

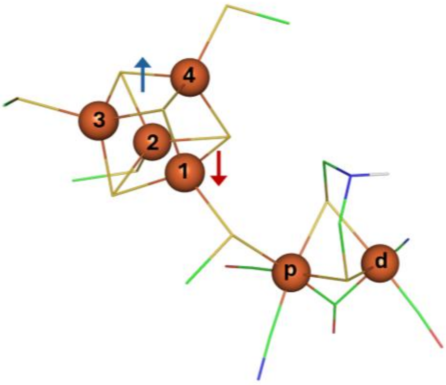


| $H_{inact}$ model/ligand identity | *Fe$_d$* | *Fe$_p$* | *Fe1* | *Fe2* | *Fe3* | *Fe4* |
|---|---|---|---|---|---|---|
| *BP86-D4/def-TZVP/COSMO ε=4* | | | | | | |
| **C367-S$^-$ (Cα constrained)** | 0.00 | 0.00 | -2.94 | -3.04 | 3.15 | 3.16 |
| **C367-S$^-$ (Cα relaxed)** | 0.00 | 0.00 | -2.94 | -3.03 | 3.15 | 3.16 |
| *TPSS-D4/def-TZVP/COSMO ε=4* | | | | | | |
| **C367-S$^-$ (Cα constrained)** | 0.00 | 0.00 | -3.06 | -3.15 | 3.26 | 3.27 |
| **C367-S$^-$ (Cα relaxed)** | 0.00 | 0.00 | -3.07 | -3.15 | 3.27 | 3.27 |
| *TPSSh-D4/def-TZVP/COSMO ε=4* | | | | | | |
| **C367-S$^-$ (Cα constrained)** | -0.01 | -0.01 | -3.37 | -3.45 | 3.58 | 3.57 |
| **C367-S$^-$ (Cα relaxed)** | 0.01 | 0.01 | -3.40 | -3.42 | 3.63 | 3.51 |
| *B3LYP-D4/def-TZVP/COSMO ε=4* | | | | | | |
| **C367-S$^-$ (Cα constrained)** | -0.01 | -0.01 | -3.49 | -3.59 | 3.71 | 3.69 |
| **C367-S$^-$ (Cα relaxed)** | 0.01 | 0.01 | -3.56 | -3.57 | 3.87 | 3.62 |

Spin densities on the Fe centers are reported for the best candidate structure of the $H_{inact}$ state across the different functionals. These values confirm the overall singlet ground state (S = 0) and the broken-symmetry solution adopted, corresponding to the inter-cubane/diiron face coupling scheme commonly used in previous studies and illustrated in the sketch below.

.

**References**


[1] Winkler, M.; Duan, J.; Rutz, A.; Felbek, C.; Scholtysek, L.; Lampret, O.; Jaenecke, J.; Apfel, U.-P.; Gilardi, G.; and Valetti, F. (2021) A safety cap protects hydrogenase from oxygen attack, *Nature communications 12*, 756.

[2] Kabsch, W. (2010) XDS, *Acta Crystallogr D Biol Crystallogr 66*, 125-132.

[3] Maly, M.; Diederichs, K.; Dohnalek, J.; and Kolenko, P. (2020) Paired refinement under the control of PAIREF, *IUCrJ 7*, 681-692.

[4] Maly, M.; Diederichs, K.; Dohnalek, J.; and Kolenko, P. (2021) PAIREF: paired refinement also for Phenix users, *Acta Crystallogr F Struct Biol Commun 77*, 226-229.

[5] Duan, J.; Rutz, A.; Kawamoto, A.; Naskar, S.; Edenharter, K.; Leimkuhler, S.; Hofmann, E.; Happe, T.; and Kurisu, G. (2025) Structural determinants of oxygen resistance and $Zn^{2+}$-mediated stability of the [FeFe]-hydrogenase from Clostridium beijerinckii, *Proc Natl Acad Sci U S A 122*, e2416233122.

[6] Afonine, P. V.; Grosse-Kunstleve, R. W.; Echols, N.; Headd, J. J.; Moriarty, N. W.; Mustyakimov, M.; Terwilliger, T. C.; Urzhumtsev, A.; Zwart, P. H.; and Adams, P. D. (2012) Towards automated crystallographic structure refinement with phenix. refine, *Biological crystallography 68*, 352-367.

[7] Emsley, P.; Lohkamp, B.; Scott, W. G.; and Cowtan, K. (2010) Features and development of Coot, *Biological crystallography 66*, 486-501.

[8] Artz, J. H.; Zadvornyy, O. A.; Mulder, D. W.; Keable, S. M.; Cohen, A. E.; Ratzloff, M. W.; Williams, S. G.; Ginovska, B.; Kumar, N.; Song, J.; McPhillips, S. E.; Davidson, C. M.; Lyubimov, A. Y.; Pence, N.; Schut, G. J.; Jones, A. K.; Soltis, S. M.; Adams, M. W. W.; Raugei, S.; King, P. W.; and Peters, J. W. (2020) Tuning Catalytic Bias of Hydrogen Gas Producing Hydrogenases, *J. Am. Chem. Soc. 142*, 1227-1235.

[9] Duan, J.; Hemschemeier, A.; Burr, D. J.; Stripp, S. T.; Hofmann, E.; and Happe, T. (2023) Cyanide Binding to [FeFe]-Hydrogenase Stabilizes the Alternative Configuration of the Proton Transfer Pathway, *Angewandte Chemie International Edition 62*, e202216903.

[10] Franzke, Y. J.; Holzer, C.; Andersen, J. H.; Begusic, T.; Bruder, F.; Coriani, S.; Della Sala, F.; Fabiano, E.; Fedotov, D. A.; and Fürst, S. (2023) TURBOMOLE: Today and tomorrow, *Journal of chemical theory computation 19*, 6859-6890.

[11] Schäfer, A.; Huber, C.; and Ahlrichs, R. (1994) Fully optimized contracted Gaussian basis sets of triple zeta valence quality for atoms Li to Kr, *The Journal of chemical physics 100*, 5829-5835.

[12] Perdew, J. P. (1986) Density-functional approximation for the correlation energy of the inhomogeneous electron gas, *Physical review B 33*, 8822.

[13] Becke, A. D. (1988) Density-functional exchange-energy approximation with correct asymptotic behavior, *Physical review A 38*, 3098.

[14] Lee, C.; Yang, W.; and Parr, R. G. (1988) Development of the Colle-Salvetti correlation-energy formula into a functional of the electron density, *Physical review B 37*, 785.

[15] Becke, A. D. (1993) Density-functional thermochemistry. III. The role of exact exchange, *The Journal of chemical physics 98*, 5648-5652.

[16] Tao, J.; Perdew, J. P.; Staroverov, V. N.; and Scuseria, G. E. (2003) Climbing the density functional ladder: Nonempirical meta–generalized

gradient approximation designed for molecules and solids, *Physical review letters 91*, 146401.
[17] Staroverov, V. N.; Scuseria, G. E.; Tao, J.; and Perdew, J. P. (2003) Comparative assessment of a new nonempirical density functional: Molecules and hydrogen-bonded complexes, *Physical review B 119*, 12129-12137.
[18] Caldeweyher, E.; Ehlert, S.; Hansen, A.; Neugebauer, H.; Spicher, S.; Bannwarth, C.; and Grimme, S. (2019) A generally applicable atomic-charge dependent London dispersion correction, *The Journal of chemical physics 150*.
[19] Schäfer, A.; Klamt, A.; Sattel, D.; Lohrenz, J. C.; and Eckert, F. (2000) COSMO Implementation in TURBOMOLE: Extension of an efficient quantum chemical code towards liquid systems, *Physical Chemistry Chemical Physics 2*, 2187-2193.
[20] Kubas, A.; De Sancho, D.; Best, R. B.; and Blumberger, J. (2014) Aerobic Damage to [FeFe]-Hydrogenases: Activation Barriers for the Chemical Attachment of O2, *Angewandte Chemie International Edition 53*, 4081-4084.
[21] Felbek, C.; Arrigoni, F.; de Sancho, D.; Jacq-Bailly, A.; Best, R. B.; Fourmond, V.; Bertini, L.; and Léger, C. (2021) Mechanism of hydrogen sulfide-dependent inhibition of FeFe hydrogenase, *ACS Catalysis 11*, 15162-15176.
[22] Just, G. H.; Lefebvre, C.; Rajamani, A.; Khartabil, H.; Pilmé, J.; and Hénon, É. (2026) A visualizable and widely applicable steric repulsion descriptor for guiding experimental chemistry, *Chemical Science*.
[23] Lefebvre, C.; Klein, J.; Khartabil, H.; Boisson, J. C.; and Hénon, E. (2023) IGMPlot: A program to identify, characterize, and quantify molecular interactions, *Journal of Computational Chemistry 44*, 1750-1766.
